# Elucidating molecular responses to spittlebug attack in *Paspalum regnellii*

**DOI:** 10.1101/2023.08.29.555367

**Authors:** Isabela dos Santos Begnami, Alexandre Hild Aono, Diego da Silva Graciano, Sandra Maria Carmello-Guerreiro, Rebecca Caroline Ulbricht Ferreira, Wilson Malagó, Marcos Rafael Gusmão, Anete Pereira de Souza, Bianca Baccili Zanotto Vigna

**Author notes:** CORRESPONDING AUTHOR Drª. Bianca Baccili Zanotto Vigna Embrapa Pecuária Sudeste. São Carlos, São Paulo, Brazil. CEP 13560-970.

## Abstract

Spittlebugs cause large production losses that affect agribusiness worldwide. Understanding plant-herbivore interactions at the molecular level may be the key to developing resistant cultivars. After a nymph survival experiment, root transcriptomes were assembled from two *Paspalum regnellii* genotypes (BGP 248 and 344) with different first-line defense strategies, with no infestation and at two times after the initial attack of the spittlebug (*Mahanarva spectabilis)* nymph, thus integrating differential expression analysis and biological network modeling supplemented by root anatomical analysis. Gene Ontology terms related to different stress responses, such as salicylic acid catabolic process, were enriched in BGP 248, while some specific to spittlebugs, such as response to herbivores, were enriched in BGP 344. Enriched pathways were related to structural differences between genotypes, such as those related to cutin, suberin and wax biosynthesis. BGP 344 also presented pathways related to induced defense, such as glutathione metabolism. Metabolic networks highlighted kinases, and coexpression networks demonstrated a complex cascade response that included lncRNAs. This study provides the first molecular insights into the defense mechanisms of *P. regnellii* against *M. spectabilis.* The genotype with the fastest response to insect attack and highest nymph mortality (BGP 344) presented kinases and an enriched glutathione pathway, in addition to constitutive barriers, such as lignin, which can make it difficult for the insect to colonize the plant.

## 1 INTRODUCTION

Livestock are responsible for ensuring the livelihoods of approximately 1.3 billion people worldwide. In addition, pasture areas total more than half of the planet’s land surface for cattle raising or animal feed cultivation (FAO). Brazil is one of the largest producers, exporters, and consumers of beef in the world. In 2022, approximately 194 million cattle were distributed on more than 160 million hectares and accounted for more than US$ 180 billion in all businesses and movements, including inputs and animal nutrition, which consists of forage (ABIEC, 2022).

The main agricultural pest of tropical pastures is spittlebugs (Hemiptera: Cercopidae), which cause global economic losses of more than US$ 2 billion a year (Thompson, 2004) because of quantitative and qualitative damage that reduces dry matter volume and decreases nutritional capacity (Congio et al., 2012). In Brazil, the genus *Mahanarva* became highly reported as an agricultural pest of forage grasses and sugarcane in the 1990s (Congio et al., 2020), with *Mahanarva spectabilis* (Distant, 1909) being responsible for limiting livestock production of beef and milk (Auad et al., 2007).

In the case of spittlebug infestation, the nymph settles in the roots and feeds on the xylem, damaging the vessels and preventing full nutrient transport, while the adults attack the leaves, breaking the cellular wall and reducing the amount of chlorophyll until it leads to tissue death (Alvarenga et al., 2019). The plant defense strategies used to combat herbivores generally involve constitutive factors that are always present, such as morphological characteristics and chemically induced defenses (Walters, 2011). These secondary metabolites are important for recognizing and responding to environmental stressors, such as herbivory. They are not synthesized all the time due to high energy expenditure, but they start to be produced when the plant detects an attack, allowing plant survival and preventing insects from developing (Singh et al., 2021).

The chemical control of insects, which are common in agriculture worldwide, is ecologically and economically unfeasible in the case of spittlebugs due to the large area of pasture. As a result, the best management method involves the development of resistant forage cultivars (Silva et al., 2019). This process traditionally involves crossing individuals to generate hybrids and fixing and establishing them in the field for testing, but it is time-consuming and does not guarantee the achievement of the desired characteristic without other trait loss (Lenaerts et al., 2019).

An alternative is to use a native genus that is more resistant than other forages to spittlebugs, such as *Paspalum* (Gusmão et al., 2016), which originated from the tropical and subtropical regions of the Americas and is composed of species with high potential for pasture (Acuña et al., 2019; Novo et al., 2016). Embrapa Pecuária Sudeste maintains a *Paspalum* germplasm bank that allows breeding program development using *Paspalum regnellii* as a source of sexuality in crosses (Matta et al., 2023). Moreover, it is a perennial species with good biomass production, some resistance to shading and high forage potential (Bortolin et al., 2019).

Currently, genomic information is essential in breeding programs to increase gains and reduce the amount of time needed, but the application of these tools has been very limited (Pereira et al., 2018). *P. regnellii*, for example, has only a few nucleotide sequences publicly available in NCBI and no published genetic studies. Therefore, transcriptome analysis is an alternative method that can provide complete transcript expression profiles under specific response conditions (Egan et al., 2012), including new and rare transcripts even for species without prior genomic information (Wang et al., 2009). This analysis has been successfully applied in works that aimed to identify pest resistance genes in different plants, including species from the Poaceae family (Zhao et al., 2017), and has generated significant differential expression results related to responses in infested plants (Becker et al., 2017; Wang et al., 2018).

In addition to the lack of genomic studies on the species that are the subject of this study, no expression analysis of resistance to spittlebugs has been reported until now, representing an important information gap for such a complex process. In this context, this research aimed to characterize the roots of two *P. regnellii* genotypes with differential responses to *M. spectabilis* attack in terms of initial nymph mortality, root anatomy, transcriptomic analysis and coexpression and metabolic networks to elucidate the genetic and molecular mechanisms involved in plant defense.

## 2 MATERIALS AND METHODS

### 2.1 Resistance experiment

Bioassays were conducted to determine the resistance level of *P. regnellii* accessions to *M. spectabilis* in a greenhouse at Embrapa Pecuária Sudeste, São Carlos, SP, Brazil, at 21°96’17” S and 47°84’21” W, at a mean altitude of 856 m. The effects on genotypes were evaluated using the nymphal survival parameter to test the antibiosis resistance hypothesis with methodology adapted from Lapointe et al. (1992), Valério et al. (1997) and Gusmão et al. (2016). Genotypes BGP-215, BGP-248, BGP-258, BGP-341, BGP-344, BGP-345 and BGP-397 were established from seeds of plants retained in the *Paspalum* Active Germplasm Bank at Embrapa Pecuária Sudeste and compared with the susceptible controls *Urochloa brizantha* (Hochst. ex A. Rich.) R. D. Webster cv. Marandu and *U. decumbens* (Stapf) R. D. Webster cv. Basilisk.

The seeds were sown in a 162-cell JKS® model tray (50 cm³) filled with substrate based on sphagnum peat, expanded vermiculite, dolomitic limestone, agricultural plaster and NPK fertilizer (Carolina®). After 30 days, the seedlings were transplanted to 480 ml Styrofoam cups (DART® 480J32) containing the substrate for Vivatto® Plus plants. Each one was covered with J32 DART® Styrofoam cups containing a central hole for the plant aerial part to maintain the absence of light on the lap and allow superficial root growth to ensure local feed for nymphs hatched.

The experimental design was completely randomized, with ten replicates. Plants were infested with insect eggs 30 days after transplanting using five eggs obtained according to Auad et al. (2007). At the time of plant infestation, the eggs showed complete embryonic development, that is, the operculum and reddish ocular and glandular spots appeared (Valério, 2009). The nymphal survival evaluations were performed weekly, from plant infestation with eggs until the adult insects emerged.

Finally, the nymph survival percentage was calculated for each plant genotype and transformed into SQUARE ROOT (X + 0.5) and subjected to variance analysis, and the variable averages were compared by Tukey’s test (p<0.05) based on the procedures of SAS and PROC-GLM (SAS Institute, 2010).

### 2.2 RNA-seq experimental design and plant material

The experiment was carried out in a greenhouse at Embrapa Pecuária Sudeste in the 2019/2020 season. Two sexual tetraploid genotypes (BGP 248 and BGP 344) of *P. regnellii* (2n=4x=40) were selected based on the contrast identified in previous results of spittlebug resistance evaluation in different genotypes of the species.

Seedlings of the two genotypes were obtained from tillers of the same adult plant that was previously established to ensure genetic identity, and the plants were cultivated in Styrofoam cups for six weeks; therefore, the root system was well developed. Adult *M. spectabilis* insects were collected from pasture areas of *U. brizantha* cv. Marandu, and the eggs were obtained in the laboratory and incubated in a biological oxygen demand (BOD) chamber (T 25°C, RH 60% and photophase of 12 hours) until the complete development of the embryonic phase. Ten completely developed eggs were confined to the root system of the plant through an aperture made on the side of the Styrofoam cup. After infestation, the system was closed with tape to guarantee nymph fixation. The control samples were subjected to the same procedure but without infestation with eggs to simulate the same stress on the plants.

Root samples were collected in triplicate for both genotypes under three initial infestation treatments: without egg infestation (TC), after egg infestation and nymph eclosion (T1: 48 h after TC) and with nymph colonization on the roots (T2: 72 h after TC), totaling nine samples per genotype.

### 2.3 RNA sequencing, data quality and filtering

Total RNA (totRNA) from the roots was extracted with an RNeasy Plant Mini Kit (Qiagen Inc., Valencia, CA) according to the manufacturer’s protocol. A NanoDrop ND1000 spectrophotometer (Thermo Fisher Scientific Inc., Waltham, MA, USA) was used for RNA quantification. Sample efficiency and integrity were evaluated using a Bioanalyzer 2100 (Agilent Technologies, Santa Clara, CA, USA). mRNA sequencing libraries were constructed for the 18 samples and sequenced on an Illumina HiSeq 2500 v4 2×100 bp platform.

FastQC version 0.11.9 software (Babraham Bioinformatics) was used to assess the quality of the sequenced libraries. The reads were filtered with Trimmomatic version 0.39 (Bolger et al., 2014) to remove the adapters with the Illumina Adapters parameter and SlidingWindow 4:20. In addition, the software SortMeRNA version 2.1b (Kopylova et al., 2012) was used to eliminate ribosomal RNA (rRNA) with default parameters.

### 2.4 Transcriptome assembly

Since *P. regnellii* does not have a publicly available reference genome, a *de novo* assembly was performed using Trinity version 2.5.1 (Grabherr et al., 2011) with the target K-mer coverage set to 30 (normalize_max_read_cov 30) and other parameters set to the defaults. The longest isoform (unigenes) for each transcript was selected with its own software.

An assembly using the genome sequence of diploid *Paspalum notatum* (available at NCBI Genome under accession ASM2253091v1) as a reference was also tested. STAR version 2.7.3a (Dobin et al., 2013) was used to align the reads to the genome, and StringTie version 2.1.6 (Kovaka et al., 2019) was subsequently used to assemble the transcriptome.

Assembly quality was analyzed using the *Viridiplantae* dataset of BUSCO version 5.2.2 (Simão et al., 2015), which compares the sequences of the transcripts with a set of orthologous genes conserved in plants, and Bowtie2 version 2.3.3.1 (Langmead and Salzberg, 2012), which maps the reads in the transcriptome to determine the transcript abundance with default parameters.

### 2.5 Differential gene expression analysis

Salmon software version 0.14.1 (Patro et al., 2017) was used with default parameters to quantify the expression of the longest isoform. Differential gene expression (DGE) analysis was performed using the edgeR package version 3.38.4 (Robinson et al., 2010) implemented in R (R Development Core Team, 2011). Raw count data were first normalized with at least 10 counts per million (CPM) and then in a minimum of three samples. For each differential gene expression test we used a false discovery rate (FDR) cutoff ≤0.05 and a minimum log2-*fold change* (logFC) of 2.

For the analysis, samples were coded as 248 (BGP 248) or 344 (BGP 344) followed by C (control without infestation), 1 (48 h after infestation) or 2 (72 h after infestation). Tests for identifying differentially expressed genes (DEGs) were performed by comparing time points for each genotype, for example, BGP 248 in the control treatment and after 48 h of infestation (coded as 248_C vs. 248_1), and between different genotypes at the same time points, such as BGP 248 in the control treatment and BGP 344 in the control treatment (coded as 248_C vs. 344_C). A total of nine comparisons were performed, following the same codifications cited before: 248_C vs. 248_1, 248_C vs. 248_2, 248_1 vs. 248_2, 344_C vs. 344_1, 344_C vs. 344_2, 344_1 vs. 344_2, 248_C vs. 344_C, 248_1 vs. 344_1 and 248_2 vs. 344_2. As there is a biotic stressor in the plants, a principal component analysis (PCA) was performed, and a hierarchical cluster based on the Euclidean distance and grouped with the weighted pair group method with arithmetic mean (WPGMA) was constructed to confirm the similarity between the replicates.

### 2.6 Functional annotation and enrichment

Trinotate software version 3.2.1 (Bryant et al., 2017) was used to predict the open reading frames (ORFs) and translate the coding sequences to peptides. Transcript functional annotation was performed using Diamond software version 2.0.14.152 (Buchfink et al., 2021) against the SwissProt database (UniProt - https://www.uniprot.org/) with at least 60% similarity. Moreover, protein domains were found using HMMER software version 3.3.2 (Mistry et al., 2013), and finally, all the results were integrated into the Trinotate extract reports.

Gene Ontology (GO) (Ashburner et al., 2000) terms were retrieved from the sequences that were functionally annotated in the database. We restricted the results for plants (*Arabidopsis thaliana*) and enriched the DEGs in each contrast with the topGO R package (Alexa and Rahnenfuhrer, 2022) for biological processes using Fisher’s exact test (*p* value < 0.01). The enriched terms were submitted to REVIGO (Supek et al., 2011) with medium similarity allowed (0.7) for data summarization.

Additionally, the Kyoto Encyclopedia of Genes and Genomes (KEGG) (Kanehisa and Goto, 2000) database was used to map all DEGs with the *Panicum hallii* reference. Fisher’s test (*p* value < 0.01) was performed to identify enriched metabolic pathways.

### 2.7 Gene coexpression and metabolic network analysis

For the creation of gene coexpression networks for each genotype, we employed the highest reciprocal rank (HRR) methodology proposed by Mutwil et al. (2010). Initially, we computed pairwise correlations between genes using the R Pearson correlation coefficient. To ensure robust associations, we set a minimum absolute correlation coefficient threshold of 0.8 and considered a maximum of the 30 strongest edges for the network connections. The network construction process was performed using R statistical software (R Development Core Team, 2011) and igraph library version 1.3.5 (Csardi and Nepusz, 2006). Subsequently, we calculated the hub score for each gene using Kleinberg’s hub centrality algorithm (Kleinberg, 1999).

To establish a metabolic network incorporating pathways associated with the DEGs, we identified enzyme commission (EC) numbers associated with the annotations of all DEGs. Leveraging this information, we used the KEGG database (Kanehisa and Goto, 2000) to select the metabolic pathways in which these enzymes are involved. Subsequently, a unified metabolic network was constructed using BioPython version 1.78 (Cock et al., 2009), and centrality measures were evaluated using Cytoscape version 3.91.1 (Shannon et al., 2003).

### 2.8 Quantitative reverse transcription PCR (RT‒qPCR)

The DEGs were validated by RT‒qPCR. Ten target genes were selected from the DEG list by the fold-change value. In addition, five housekeeping genes were chosen based on the literature (Andrade et al., 2017; Gimeno et al., 2014; Liu et al., 2017; Takamori et al., 2017). Primer pairs (Table S1) were designed using Primer3Input (Untergasser et al., 2012). Dimers, heterodimers and hairpins were checked in NetPrimer (Premier Biosoft).

cDNA synthesis was performed with the GoScript™ Reverse Transcription System A5000 (Promega, Madison, WI, USA) according to the manufacturer’s protocol. RT‒qPCR was performed on a QuantStudio 6 Pro (Applied Biosystems, Waltham, MA, USA) using GoTaq® qPCR Master MixA6001 (Promega, Madison, WI, USA) with the following protocol: 95°C for 120 s, followed by 40 cycles of 95°C for 15 s and 60 s at 60°C. Melting curves were obtained by heating from 60 to 95°C to verify the products after amplification. All qPCR experiments were performed using two technical and three biological replicates and analyzed with Design and Analysis Software 2.6.0 (Thermo Fisher Scientific Inc., Waltham, MA, USA).

### 2.9 Root anatomical analysis

As in the RNA-seq experiment, the seedlings were obtained from tillers of the same plant and cultivated for six weeks for the root system to develop. Roots of both genotypes, without spittlebug infestation, were fixed in BNF (phosphate buffer solution, formalin; 9:1 v/v) for 48 hours (Lillie, 1965) and stored in 70% ethanol. The material was dehydrated in an ethanol series including hydroxyethyl-methacrylate (Historesin® Leica) following the manufacturer’s recommendations. Five-micron-thick cross-sections were made with the aid of a Microm HM 340E rotary microtome (Thermo Fisher Scientific Inc., Waltham, Massachusetts, USA).

The slides were stained with 0.05% toluidine blue in citrate buffer (pH = 4.5) (O’Brien et al., 1964) and finally mounted on Entellan® synthetic resin (Merck KGaA, Darmstadt, Germany). Images were captured with a digital camera (Olympus DP71) coupled to an Olympus BX51 optical microscope (Olympus Optical Co., Ltd., Japan). To detect the main chemical compounds present in the cell walls of root cells, we used the following histochemical tests: acidified phloroglucinol for lignin detection (Johansen, 1940) and Nile red fluorochrome for lipid substances under a TRITC filter (ex545/20 nm; em605/60 nm) (Greenspan et al., 1985).

## 3 RESULTS

### 3.1 Spittlebug resistance experiments

The nymph survival average, percentage and standard error were calculated from the seventh day of plant infestation with *M. spectabilis* eggs until the emergence of the last adult insects at 56 days after infestation (Figure 1a). The percentages in the controls, *U. brizantha* cv. Marandu and *U. decumbens* cv. Basilisk were greater than those in the *P. regnellii* genotype. There were significant differences in nymph survival between *P. regnellii* accessions until 21 days after infestation, with more intensive mortality occurring in BGP 344 (green line) and less intensive mortality occurring in BGP 248 (red line) (Figure 1a).

**Figure 1.**
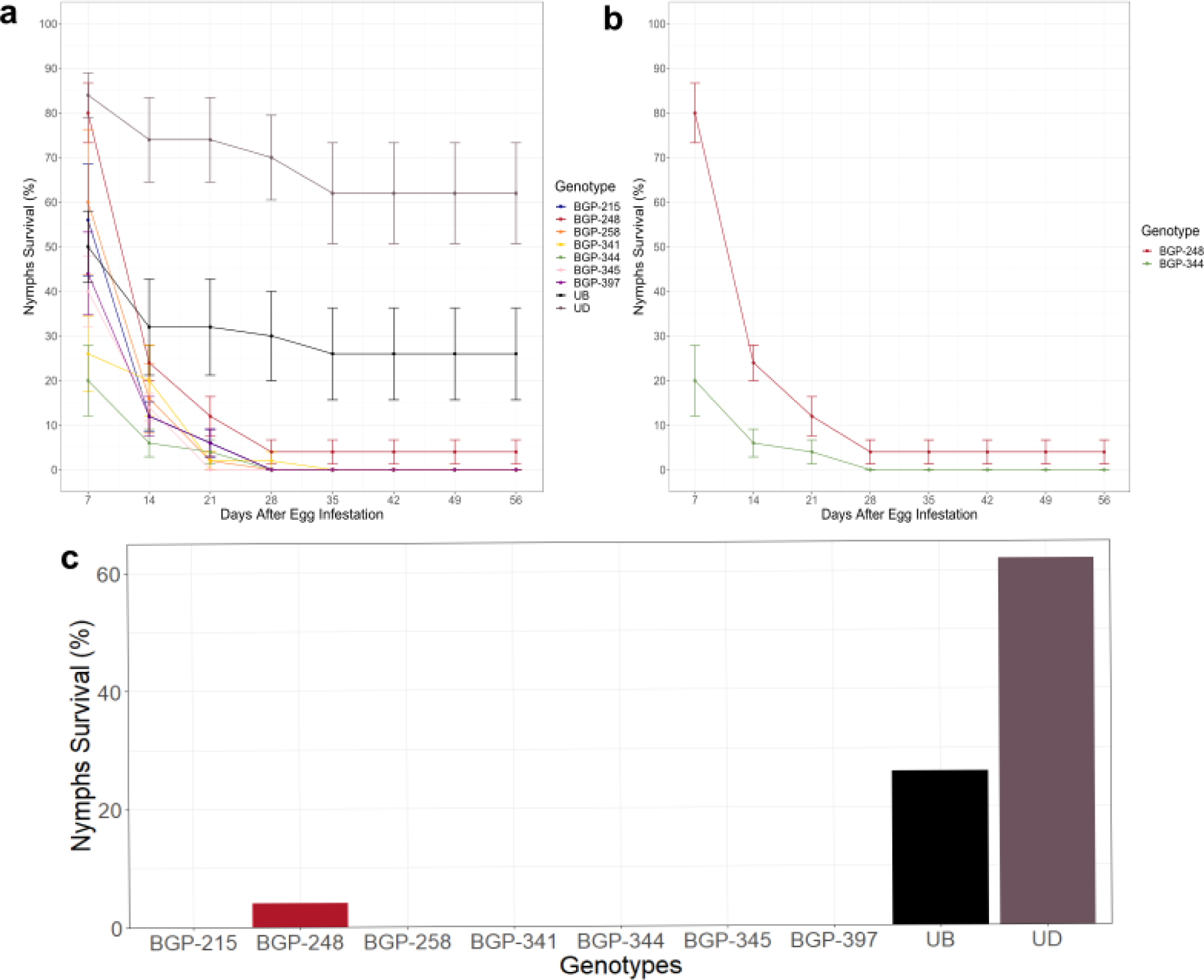
Survival percentage (average ± standard error) of *M. spectabilis* nymphs **(a)** in seven *Paspalum regnellii* genotypes in comparison with the susceptible controls *Urochloa brizantha* cv. Marandu and *U. decumbens* cv. Basilisk. **(b)** The two genotypes (BGP-248 and BGP-344) selected for transcriptome analysis based on their contrasting effects on *M. spectabilis* nymph survival. (c) Survival percentage of *M. spectabilis* nymphs until the emergence of adults 56 days after plant infestation. Means followed by a different set of letters are significantly different according to the Scheffé test at 5% significance (ANOVA: p < 0.0001; C. V = 83.42).

Significant differences were observed in the average nymph survival percentage for each *P. regnellii* genotype after several days, highlighting the difference in nymph mortality between BGP 344 and BGP 248, mainly for young nymphs of the 1st and 2nd instars, until the 21st day after infestation (Figure 1b). This is possibly related to the different strategies adopted by the genotypes in the first line of defense.

### 3.2 Transcriptome assembly and annotation

Sequencing from 18 samples from two genotypes at three different time points generated a total of 806,942,117 paired-end reads from all 18 *P. regnellii* root transcriptomes sampled. FastQC software revealed that all 18 samples presented a similar quality profile, ∼19.97% of the reads were discarded as low-quality sequences (the remaining 645,827,820 reads), and ∼4.95% were discarded as ribosomal RNA residues (the remaining 613,888,166 reads) after filtering.

Both the *de novo* assembly with only the longest isoform (unigenes) and the reference-based assembly using the *P. notatum* genome were performed, and their statistics were compared (Table 1). The second method generated good transcript results, but in the initial stage of aligning the reads with the reference, we obtained only 55.96% success, losing a large amount of data. In addition, it was possible to observe with BUSCO that the assembly generated many duplicate transcripts. The low alignment (Table 1) with another species of the same genus (*P. notatum*) suggested genomic variation even within the genus itself, showing differences in genome composition, which was also confirmed by the low number of annotated transcripts. This was expected because of the great genomic complexity and diversity present in *Paspalum*, which resulted in the separation of the two species into botanical informal groups primarily by Chase (1929), phylogenetic segregation in distinct clades (Delfini et al., 2023) and different genomic compositions by comparing a diploid genome from the Notata group (NN) and an allotetraploid from the Virgata group (IIJJ) (Cidade et al., 2013).

**Table 1.**
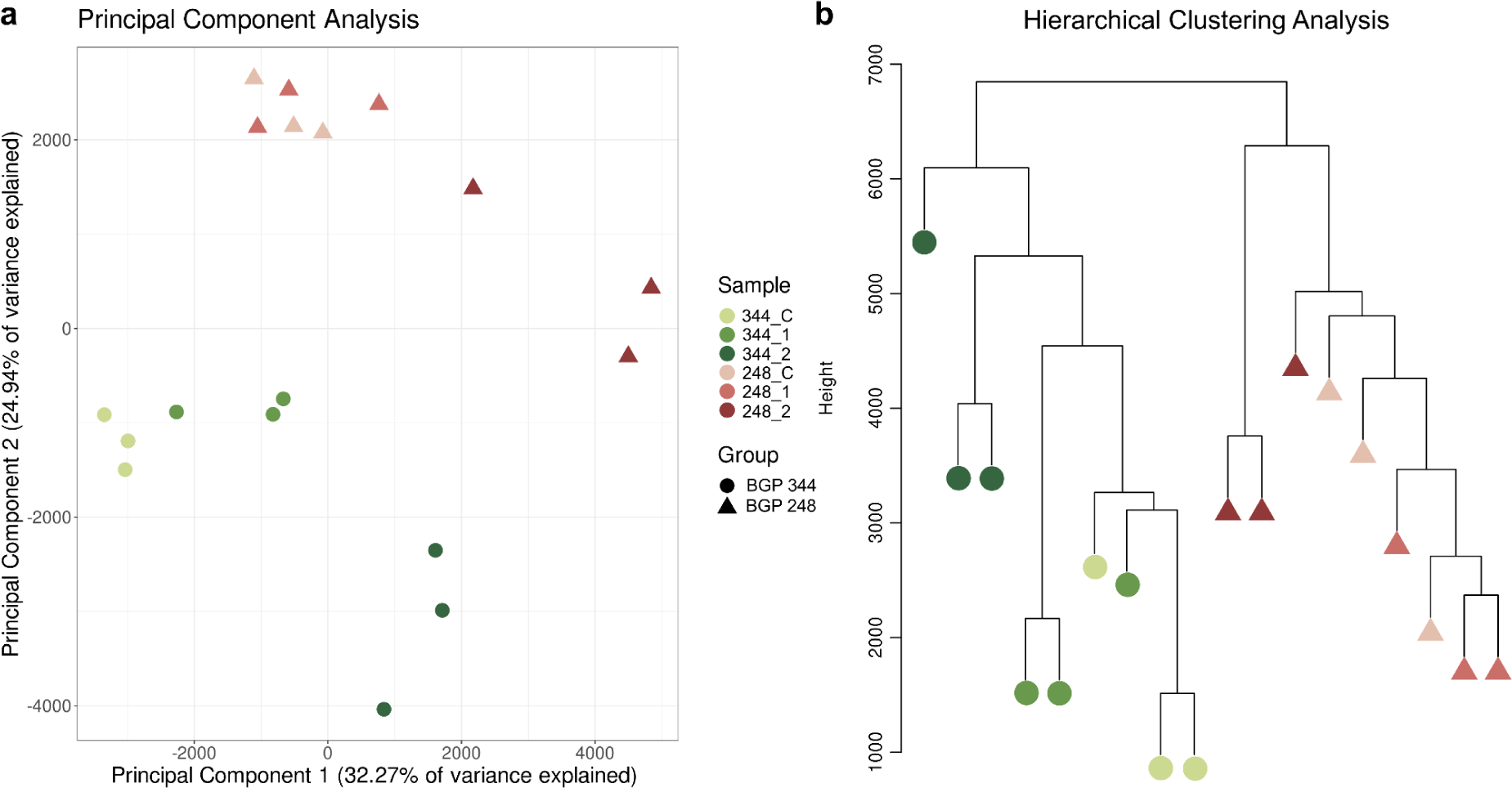
Comparison between the *de novo* assembly (Trinity - Unigenes) and the genome reference (StringTie) followed by their quality statistics.

Despite the large transcript number and lower values of N50 and mean size, transcriptome *de novo* assembly in polyploid plants is naturally complex because of the homeologous presence that creates many transcripts with structural abnormalities (Payá-Milans et al., 2018). Trinity software lost less data and generated successful results for other *Paspalum* species (de Oliveira et al., 2020); therefore, this assembly was selected for subsequent analyses. Almost 16% (92,035) of the transcripts were annotated against the SwissProt database, and 7.3% (42,040) of the proteins were predicted with Transdecoder.

### 3.3 Differential expression analysis and enrichment

After CPM normalization and expression filtering of the raw count data for all samples, we obtained a total of 21,508 genes, which were used as the data universe for the next analysis. In both the PCA and WPGMA results (Figure 2), it was possible to observe a clear genotype separation in addition to a proximity between the samples of the control treatment and 48 h after plant infestation, while the last treatment was more distant.

**Figure 2.**
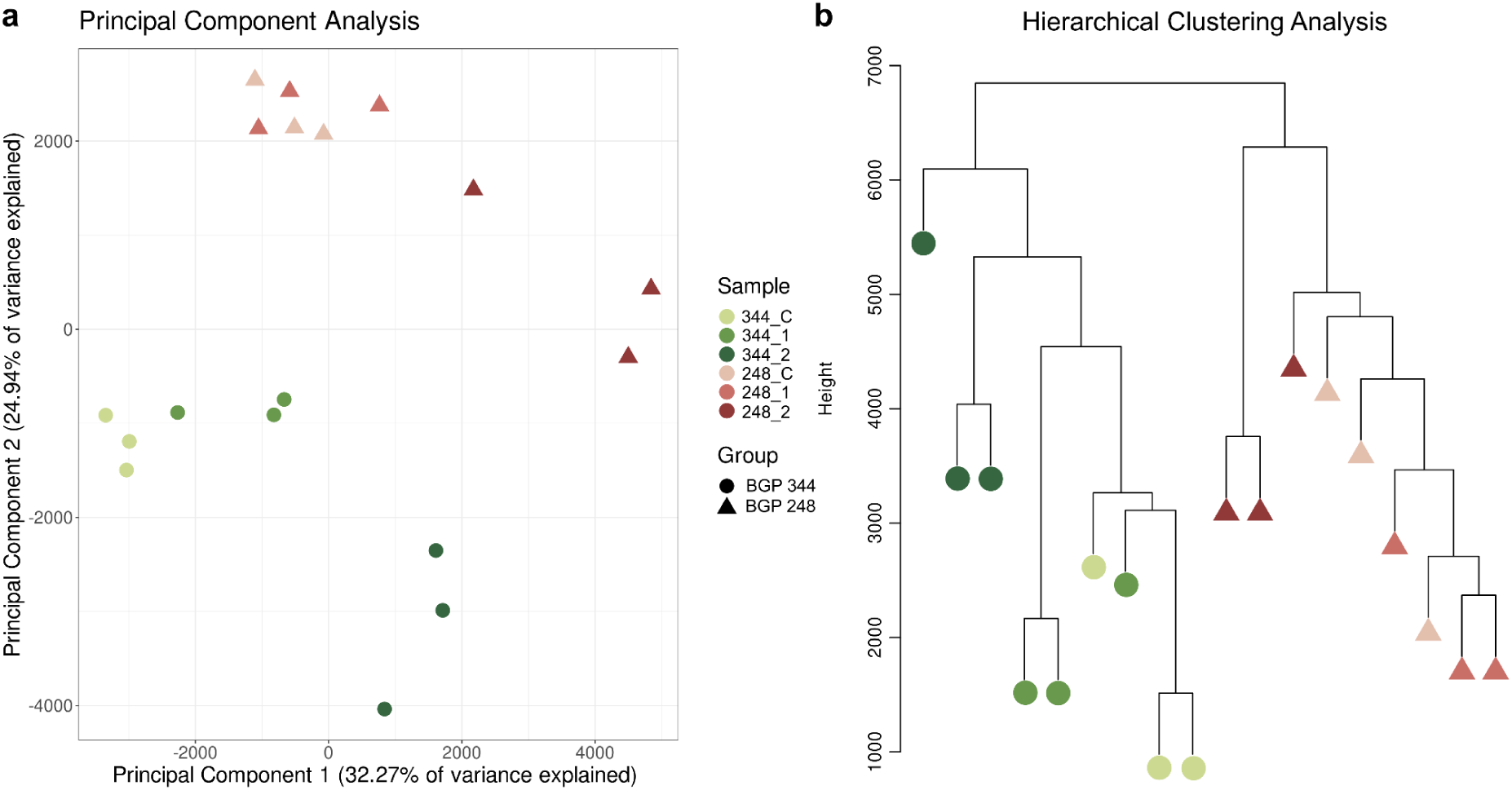
Sample distribution by **(a)** principal component analysis. **(b)** Hierarchical clustering analysis was performed with Euclidean distance and the weighted pair group method with arithmetic mean (WPGMA).

The genes were subjected to differential expression analysis (Table S1), and contrasts between genotypes at the same treatment time revealed many DEGs even in the control, showing a considerable basal difference (Table S2). In the treatment comparison, neither genotype showed DEGs between TC and T1, but the expression of many genes changed after 72 h of infestation (T2), mostly in the BGP 344 genotype.

Venn diagrams were created to investigate the intersection of DEGs between the comparisons (Figure 3). In contrast, in the same genotype, only some genes participated in more than one treatment, and most genes participated after 72 hours (Figure 3a and b). On the other hand, in genotype contrasts, 616 genes were shared between all three treatments, demonstrating the existence of an important natural difference in expression among plants even with no infestation (Figure 3c).

**Figure 3.**
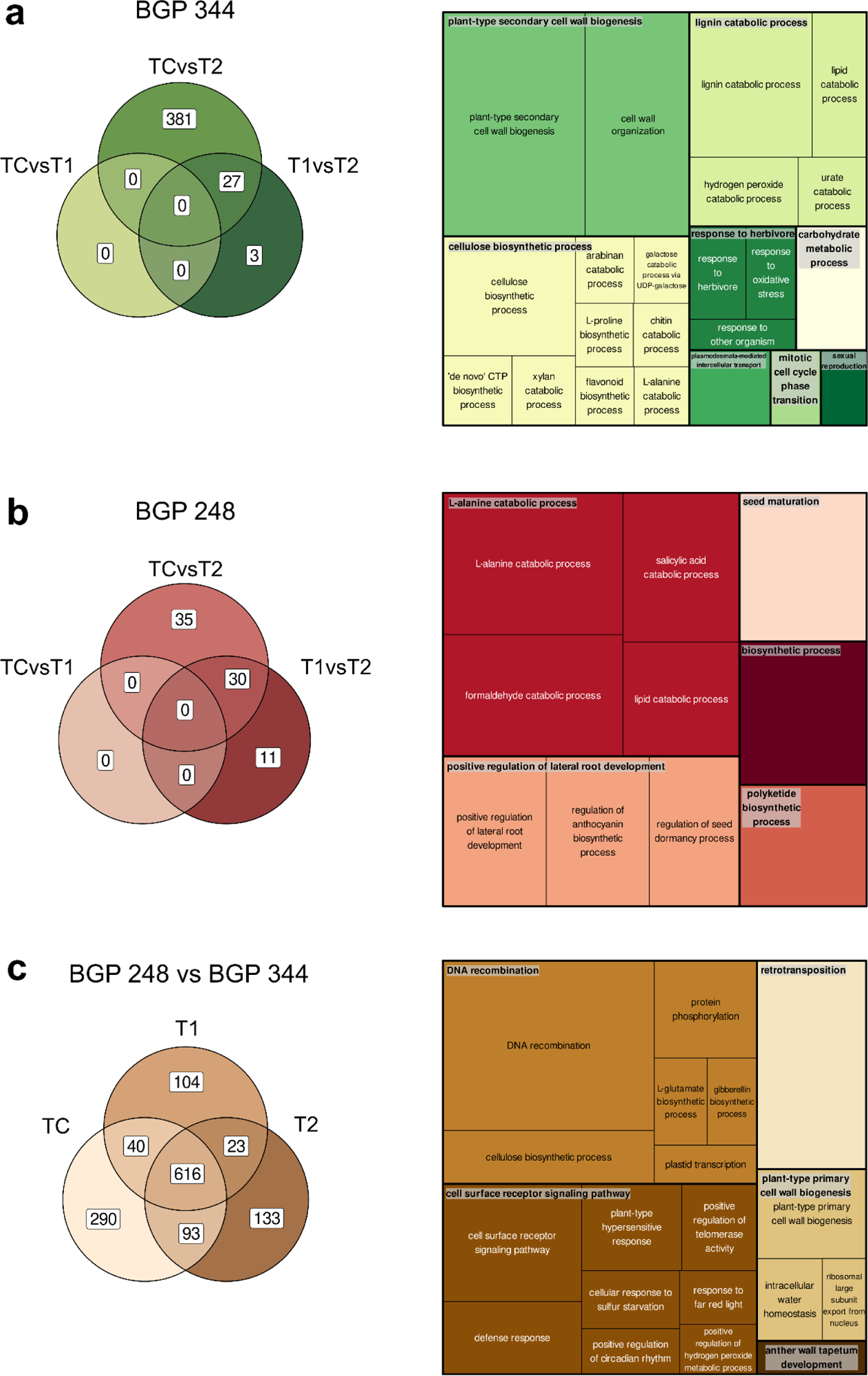
Venn diagram of DEGs and summarized Gene Ontology terms in TreeMap. **(a)** Venn diagram of all BGP 344 gene sets and GO terms enriched after 72 h (upregulated in the 344_2 subgroup). **(b)** Venn diagram for all BGP 248 comparisons and GO terms enriched after 72 h (upregulated in 248_2). **(c)** Venn diagram between genotypes and GOs in TreeMap for 344_2 vs. 248_2.

With the aim of exclusively selecting genes related to resistance, an analysis of the intersection between the genotypes at various time points was performed. Some transcripts had increased expression over time in both genotypes, but even in the control, they were still more highly expressed in BGP 344 (Table S3), and their annotations are possibly related to resistance: suppressor cell death (LSD1), resistance gene analogs (RGA5R) and one domain-containing protein (DUF4220). The expression of other transcripts increased in BGP 344, and these transcripts were naturally more abundant in BGP 248: transcription factors that regulate the secondary cell wall (NST1) and response inducers to ozone and pathogens (AtOZI1). Finally, those genes whose expression increased in the BGP 248 genotype were mostly related to abiotic stress (PLA7).

#### 3.3.1 GOs

The functional classes of all DEGs were enriched in 89 different GO terms of biological processes in topGO (Table S4). After spittlebug infestation, the BGP 344 genotype was closely related to resistance processes (Figure 3a), such as chitin catabolism (GO:0006032), response to oxidative stress (GO:0006979), response to other organisms (GO:0051707) and response to herbivores (GO:0080027).

In contrast, in BGP 248, there were several terms related to resistance (Figure 3b), including salicylic acid catabolic process (GO:0046244) and regulation of anthocyanin biosynthetic process (GO:0031540), but they were not very specific. Furthermore, some terms, such as seed maturation (GO:0010431) and positive regulation of lateral root development (GO:1901333), revealed that plants ignore infestation and continue to grow normally.

When we compared the genotypes in the last infestation treatment (Figure 3c), we obtained some interesting terms that had different expression levels, such as defense response (GO:0006952), plant-type hypersensitive response (GO:0009626), positive regulation of circadian rhythm (GO:0042753), retrotransposition (GO:0032197) and protein phosphorylation (GO:0006468).

#### 3.3.2 KEGG

Furthermore, the DEGs were enriched in 13 metabolic pathways according to the KEGG database (Table S5). Among these pathways, some were related to basal cellular mechanisms and were enriched in the comparisons of both genotypes: glycolysis/gluconeogenesis, fatty acid degradation, tyrosine metabolism, alpha-linolenic acid metabolism and pyruvate metabolism (Table S5). Enrichment of the DNA replication pathway, as it is more structural, was not selected for further study.

The other seven pathways that occurred in only one treatment comparison and are possibly related to resistance were chosen to construct a unique metabolic network: purine metabolism, caffeine metabolism, glutathione metabolism, galactose metabolism, cutin, suberin and wax biosynthesis, and circadian rhythm - plant, starch and sucrose metabolism.

### 3.4 Network analysis

A metabolic pathway was identified from the enriched KEGG pathways, which revealed 109 enzymes related to the resistance process (Table S6). The enzymes with greater influence on the network (“EdgeCount”) and in relationship with other enzymes (In and Outdegree) are highlighted in blue in Figure 4a and listed here: adenylate kinase (ec:2.7.4.3), glutathione reductase (ec1.8.1.7), guanylate kinase (ec:2.7.4.8), AMP deaminase (ec:3.5.4.6), apyrase (ec:3.6.1.5), nucleotide diphosphatase (ec:3.6.1.9), AMP phosphatase (ec:3.1.3.5), phosphoglucomutase (ec:5.4.2.2), glutathione dehydrogenase (ec:1.8.5.1) and uridyl transferase (ec:2.7.7.12).

**Figure 4.**
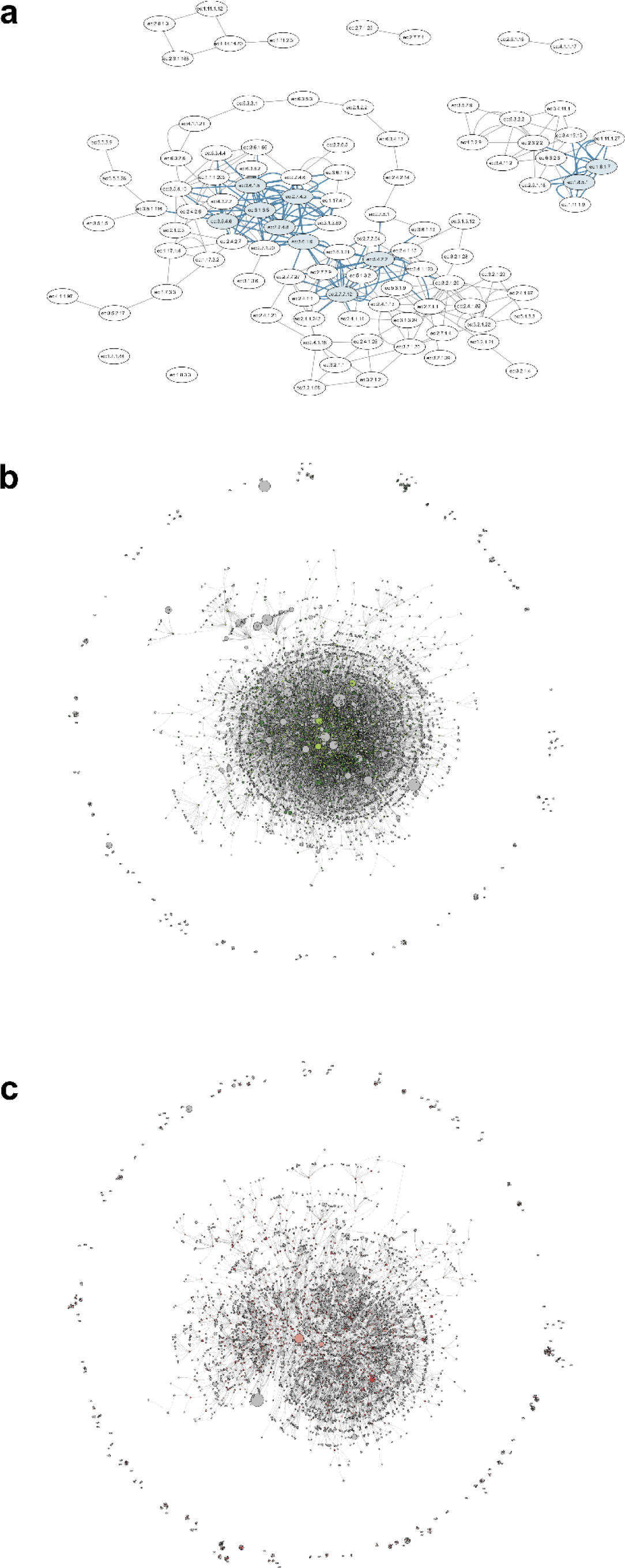
Network representations. **(a)** Metabolic network, with the most significant enzymes of enriched pathways and their connections in blue. **(b)** BGP-344 coexpression network. The green nodes represent the hub genes. **(c)** BGP-248 coexpression network. The pink nodes represent the hub genes.

Gene coexpression networks were constructed for each genotype separately. The BGP 344 network (Figure 4b) had 7,410 nodes (Table S7), so we observed many more hubs and connections in its network than did the BGP 248 network (Figure 4c), which had 5,275 nodes (Table S8). This suggests a large difference in transcript number between genotypes as well as in DEG analyses, showing the complexity of the genetic response. Furthermore, some of the transcripts with the greatest impact on the networks because of the higher “degree” number were not characterized or not annotated.

### 3.5 RT‒qPCR validation

A total of ten transcripts were selected to validate the *in silico* differential gene expression, and in two of them, flavonol 3-sulfotransferase (F3) and IQ domain-containing protein (IQ), primers did not score properly, so they were discarded. Validation was performed with two unknown transcripts and six annotated genes that were chosen because of the considerable difference in expression (Figure S1).

In addition, two of five normalizers were selected as endogenous controls in the experiment based on lowest quantification cycle (Cq) variation values using five methods collected by RefFinder (Xie et al., 202): NADH dehydrogenase ubiquinone 1 alpha (NDAU1) and splicing factor U2af small subunit B (U2AFB) (data not shown). The expression patterns generated by RT‒qPCR analysis agreed with the transcriptome data, proving the accuracy of the bioinformatic results (Figure S1a), in addition to the data showing good correlation (Figure S1b).

### 3.6 Root anatomical analysis

We performed a comparative anatomical analysis between roots of the two *P. regnellii* genotypes without biotic stress to investigate whether transcriptomic variations are evident in the fine root structure, where spittlebugs prefer to feed.

Both genotypes present a uniseriate epidermis, with a slightly thickened external periclinal cell wall and secretion (Figure 5a and b). Regarding the cortex, we observed the presence of an exodermis in both genotypes, characterized by a layer of cells with differences in the cell wall composition (Figure 5a and b). In BGP 248, the exodermis has a thickened cell wall consisting of lipid material (Figure 5c and d), and in BGP 344, the cells show lignified secondary cell wall deposition, as confirmed by the acidified phloroglucinol test (Figure 5e). Additionally, we observed that the exodermis cells of this genotype collapse and acquire irregular shapes, a process that begins close to the root apex (Figure 5b).

**Figure 5.**
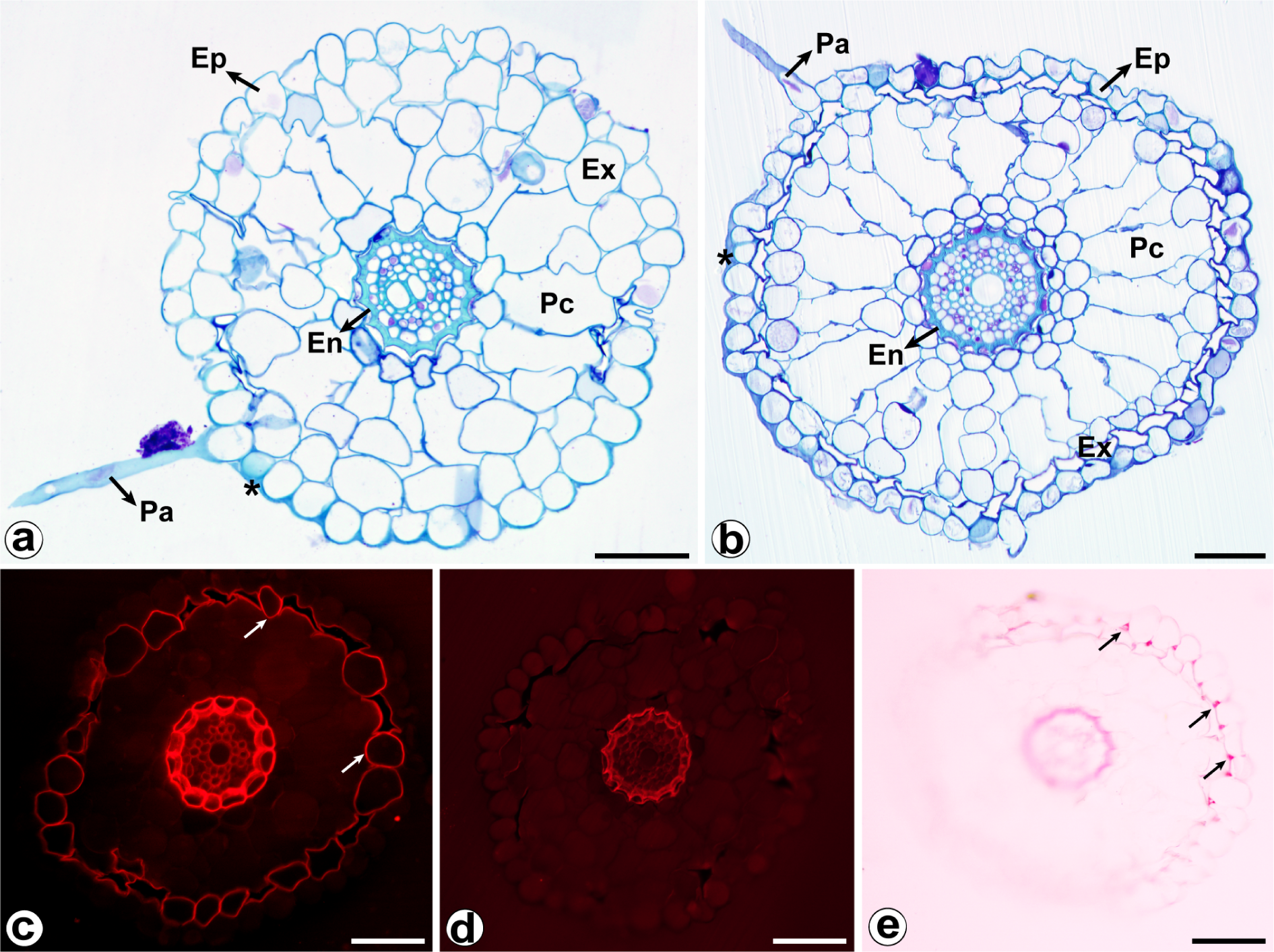
Anatomical structure and histochemical tests in root cross-sections of different *Paspalum regnellii* genotypes. **(a)** BGP 248 and **(b)** BGP 344, with asterisks indicating secretions covering the epidermis. **(c)** Positive reaction to Nile red fluorochrome, with arrows showing wall thickenings with lipid composition in BGP 248. **(d)** Negative reaction to Nile red fluorochrome on the cell walls of BGP 344 cells. **(e)** Positive reaction to acidified phloroglucinol showing lignin deposition in the cell walls of BGP 344. Abbreviations - En: endoderm; Ep: epidermis; Ex: exoderm; Rh: root hairs; Cp: cortical parenchyma. Scale bars: 50 μm (a-e).

The cortical parenchyma in both genotypes has wide intercellular spaces and forms an aerenchyma, which is more developed in BGP 344, contributing to the formation of irregular cells of the exodermis (Figure 5a and b). In the BGP 344 genotype, the internal cellular layers of the cortical parenchyma present globular cells arranged in a spiral pattern around the endodermis (Figure 5b). The endodermis has Casparian strips, and the cells exhibit “U”-shaped thickening of the lignified wall (Figure 5a and b). The pericycle is uniseriate, and the vascular system is composed of a few protoxylem poles alternating with phloem strands and a few central metaxylem elements (Figure 5a and b).

## 4 DISCUSSION

In the current scenario of plant breeding, it is customary to explore genomic regions responsible for governing traits of interest, laying the basis for marker-assisted selection and gene editing. However, resistance to herbivory presents a more intricate challenge (Dinardo-Miranda et al., 2014), characterized by a complex genetic architecture coordinated by a diverse set of genes associated with numerous quantitative trait loci (QTLs) (Zhang et al., 2020a). Differential expression analysis enables the elucidation of how plants perceive and respond to different environmental stressors, aiding in the identification of genes directly influenced by insects (Cui et al., 2017). These identified genes can subsequently serve as focal points for further exploration, including potential targets for genetic engineering (Ramstein et al., 2019). In this context, our study performed a comprehensive characterization of the molecular mechanisms triggered by spittlebug attacks, thus enhancing our understanding of the diverse strategies plants employ in response to herbivory.

Currently, the predominant pasture area in South America is composed of African species with high productivity levels, such as *Panicum* and *Urochloa* (Matias et al., 2021). This prevalence can be attributed to breeding efforts focused on enhancing field performance (Casler and Santen, 2010). However, despite their productivity, these species are susceptible to insect attacks, posing major concerns for livestock management. In Brazil, one of the most damaging pests is the spittlebug *Mahanarva spectabilis*, which heavily impacts productivity. Notably, the native genus *Paspalum* appears to suppress the performance of this pest, possibly due to coevolutionary adaptations (Gusmão et al., 2016). Despite the large economic losses caused by spittlebug attacks, investigations into the molecular mechanisms underlying such resistance remain scarce. The resistance experiment carried out in a greenhouse revealed differences between the *P. regnellii* genotypes, with an emphasis on BGP-344 and BGP-248, whose initial nymph mortality was more pronounced in BGP-344, indicating its potential as a rich source of genetic information for resistance mechanisms. The *de novo* transcriptome assembly presented in this study constitutes the first comprehensive molecular resource for elucidating the intricate interplay between forage grasses and spittlebugs.

To enhance the reliability and specificity of our data regarding resistance mechanisms, we opted to compare *P. regnellii* with a phylogenetically distant susceptible species, such as *Urochloa* (Soreng et al., 2022). The considerable number of genetic differences arising from factors unrelated to insect feeding could contradict the interpretation of these results. Instead, our approach consisted of analyzing transcripts after two and three days of infestation to delineate the preliminary responses observed between genotypes in the field experiment. Notably, we observed that BGP 344 resulted in greater initial nymph mortality than did BGP 248 for up to 21 days (Figure 1a and b). This discrepancy can be attributed to the well-established defense mechanisms of tropical grasses against insect pests and pathogens. These defenses start with physical barriers, including cell wall adaptations, to avoid colonization for feeding (Zhang et al., 2020b). Lignin, for instance, serves as a mechanical component that reduces plant palatability for herbivores (Xiao et al., 2023), increases cellular rigidity, and prevents insect colonization (Lynch et al., 2021; Wu et al., 2022), thereby conferring resistance to antixenosis (Kogan and Ortman, 1978).

Our root histochemical analyses revealed lignification exclusively in the walls of BGP 344 plants (Figure 5e), along with differentiation of the sclerenchymatous tissue into aerenchyma (Figure 5a and b). This adaptation improves gas exchange within the cellular environment under oxygen-deficient conditions, thereby mitigating hypoxia (Teixeira et al., 2022; Yamauchi and Nakazono, 2022). We believe that these anatomical predispositions make BGP 344 less susceptible to host insects than BGP 248. This assertion is corroborated by the GO analysis, which revealed enrichment in BGP 344 (Figure 3a) for the terms cell wall organization and plant-type secondary cell wall biogenesis and highlighted associations with processes related to regenerative characteristics, such as cellulose biosynthetic processes and lignin catabolic processes.

The difficulty in overcoming cellular barriers and colonizing the plant tissue explains the absence of DEGs after 48 h of infestation. With the beginning of insect feeding on plant tissues, alterations primarily manifest in the conformation of the cell membrane, triggering the production of reactive oxygen species (ROS) (Walters, 2011). Subsequently, the induced defense mechanisms are activated, involving the synthesis of secondary metabolites and herbivore-induced plant volatiles (HIPVs). These compounds not only stimulate defense within the infested plant but also serve to alert neighboring plants and attract predators. The regulation of these responses is controlled by phytohormones, notably jasmonic acid (Lin et al., 2022). Within BGP 344, GO terms associated with ROS, such as hydrogen peroxide catabolic process, response to oxidative stress, and response to herbivores, were identified (Figure 3a). Notably, within BGP 248, a significant GO term related to the catabolic process of salicylic acid (SA) was observed. SA, which is typically associated with the stress response, may undergo degradation because it may not play a prominent role in defense against spittlebugs. The presence of compounds in *M. spectabilis* saliva may hinder SA signaling, although its detection has been reported in susceptible species such as *U. decumbens* postinfection (Barros et al., 2021). Furthermore, plants often prioritize resource allocation toward defense mechanisms in response to herbivore pressure, potentially at the expense of growth and development (Divekar et al., 2022). However, the BGP 248 genotype appears to continue to invest in root growth and seed development even in the presence of herbivore infestation. In this sense, we suggest delayed recognition of the pest due to the presence of distinct defense mechanisms in this genotype.

The enriched pathways observed in the genotype contrasts within the control samples, namely, circadian rhythm - plant and cutin, suberin and wax biosynthesis, are intricately related to the inherent morphological characteristics of each genotype, conferring plant defense. The circadian rhythm, which governs plant physiological processes in response to environmental differences, offers survival advantages by regulating carbon metabolism and hormone signaling pathways (Xu et al., 2022). This regulatory mechanism is closely linked to stress responses, aiding in the stabilization of plant development (Bhattacharya et al., 2017) and facilitating the liberation of metabolites against herbivore attack (Hua, 2013), thus enhancing the plant’s innate defense against spittlebugs. On the other hand, the pathways associated with cutin, suberin and wax biosynthesis constitute the formation of cuticle structures, which serve as the primary defense layer for plants (Mahatma et al., 2021). Studies suggest that cutin is involved in plant defense (Fich et al., 2016) and that suberin plays an important role as a plant root cell barrier against adverse environmental situations (Xiao et al., 2020) and pathogens (Guo et al., 2022).

The BGP 248 genotype does not exhibit any enriched pathways unique to it, whereas BGP 344 exhibits pathways associated with the induction of hormones to respond to biotic stress, such as purine and caffeine metabolism (Ashihara et al., 2008; Wang et al., 2016), along with pathways related to energy reserves (Du et al., 2020), which provide resources for carbon consumption for bear stress (Qiu et al., 2021; Zhang et al., 2021), including the metabolism of starch and sucrose (Chen et al., 2023; Thalmann and Santelia, 2017). The increased activity of these pathways suggests their involvement in the plant defense response to spittlebug attack, facilitating energy generation and the release of secondary metabolites. Moreover, when contrasting the genotypes under infestation, an elicited response commanded by the glutathione pathway was enriched in BGP 344. This pathway is one of the major contributors to antioxidant defense, regulating ROS levels under biotic and abiotic stress conditions (Gong et al., 2018; Huang et al., 2019). This induction, along with structural differences and increased compound release from the purine and caffeine pathways, distinguishes the defense response of BGP 344 from that of BGP 248, where these pathways apparently remain inactive.

The plant resistance mechanisms against herbivory involve secondary metabolites, phytohormones, and important proteins, such as kinases and transcription factors, which have been extensively studied (Wang et al., 2020). Network analyses serve as valuable approaches to highlight important elements in this context. The metabolic network (Figure 4a) revealed the significance of two enzymes within the ascorbate-glutathione (AsA-GSH) cycle, notably, dehydroascorbate reductase (DHAR) and glutathione reductase (GR), which play important roles in handling oxidative stress. These enzymes not only confer plant tolerance to stress but also participate in regenerating AsA and GSH to maintain the redox state (Dorion et al., 2021; Hasanuzzaman et al., 2019). Furthermore, enzymes from the kinase family, which are responsible for protein phosphorylation, have emerged as key players in stress tolerance mechanisms, contributing to the maintenance of cellular homeostasis (Dzeja and Terzic, 2009; Patil and Senthil-Kumar, 2020; Sekulic et al., 2002). Our analysis highlights two enzymes as potential resistance targets: adenylate kinase, which is involved in regulating salt stress in maize (Chen et al., 2022) and in supplying substrates for aluminum resistance in *U. decumbens* roots (Arroyave et al., 2018), and guanylate kinase, whose expression changes in response to armyworm herbivory in sugarcane, thereby assisting in the resistance process (Wang et al., 2021). These findings underscore the importance of glutathione pathways and the intricate relationship between the kinase family and defense mechanisms against herbivory. Furthermore, metabolic network analysis revealed novel and promising targets for enhancing plant resistance to herbivores.

Coexpression network analysis revealed hub genes pivotal to spittlebug resistance, which were not identified in other analyses but play essential roles in this intricate process by influencing several others through connections. In the BGP 344 network (Figure 4b), we identified hubs associated with a ncRNA derived from a hypothetical gene of *Setaria viridis* known to modulate plant gene expression in response to both abiotic and biotic stresses (Hou et al., 2019; Song et al., 2021). Additionally, we identified a retrotransposon gag protein and a retrovirus-related Pol polyprotein from transposon TNT 1-94 (RPPT), which regulate the transcription levels of adjacent genes in response to environmental changes (Yang et al., 2020; Hao & Zhang, 2022). Another significant hub was putative ubiquitin-like-specific protease 1 (ULP1B), which is involved in posttranslational gene modification responsible for stress defense in eukaryotes (Pedley et al., 2018; Morrell & Sadanandom, 2019). Complementing these main hubs, we annotated the trehalose 6-phosphate synthase gene, which is associated with abiotic responses (Sarkar and Sadhukhan, 2022), and two other uncharacterized genes, highlighting gaps in our understanding of the interaction between forages and spittlebugs. Our network analyses underscore the ability of networks to unveil novel discoveries and demonstrate that ncRNAs, acting as hubs, can influence the expression of genes against herbivores, expanding our understanding beyond traditional transcribed genes.

## 5 CONCLUSIONS

The present study elucidates important molecular mechanisms underlying the defense of *P. regnellii* against *M. spectabilis* spittlebugs, revealing a highly genotype-specific process. Our findings underscore the important role of morphological traits that hinder spittlebug infection, as well as the involvement of secondary metabolites, enzymes, and noncoding RNAs in plant defense mechanisms. These insights pave the way for future genomic investigations and provide essential information for forage breeders for the targeted development of spittlebug-resistant tropical forage cultivars.

## ACKNOWLEDGMENTS

We thank the “Coordenação de Aperfeiçoamento de Pessoal de Nível Superior” (CAPES) and the Programa de “Excelência Acadêmica” (PROEX) for the ISB master’s scholarship (#88887.600436/2021-00). We thank Embrapa (Grant #20.18.01.014.00.00) and Unipasto (Association for Fostering Research on Forage Improvement) for funding and support. We thank “Fundação de Amparo à Pesquisa do Estado de São Paulo” (FAPESP) for scholarships of AHA PhD scholarship (#2019/03232-6) and RCUF PD fellowship (#2018/19219-6) and “Conselho Nacional de Desenvolvimento Científico e Tecnológico” (CNPq) for APS research grant (#312777/2018) and DSG scholarship (#140174/2021-4).

## AUTHOR CONTRIBUTIONS

MRG, WMJ, and BBZV conceived the study. ISB, DSG, MRG, WMJ, and BBZV obtained the data. ISB, AHA, and RCUF analyzed the transcriptome and network data; ISB and WMJ analyzed the RT‒qPCR results; and DSG and SMC-G analyzed the anatomy. MRG, BBZV, and APS acquired funding and oversaw the work. ISB led the writing of the manuscript, and all authors contributed to the writing. All authors revised the work critically and approved the submitted version.

## DATA AVAILABILITY STATEMENT

The data that support the findings of this study are openly available in the Sequence Read Archive (SRA) at https://www.ncbi.nlm.nih.gov/sra, reference number PRJNA999588.

## SUPPLEMENTARY FIGURES AND TABLES

**Figure S1.**
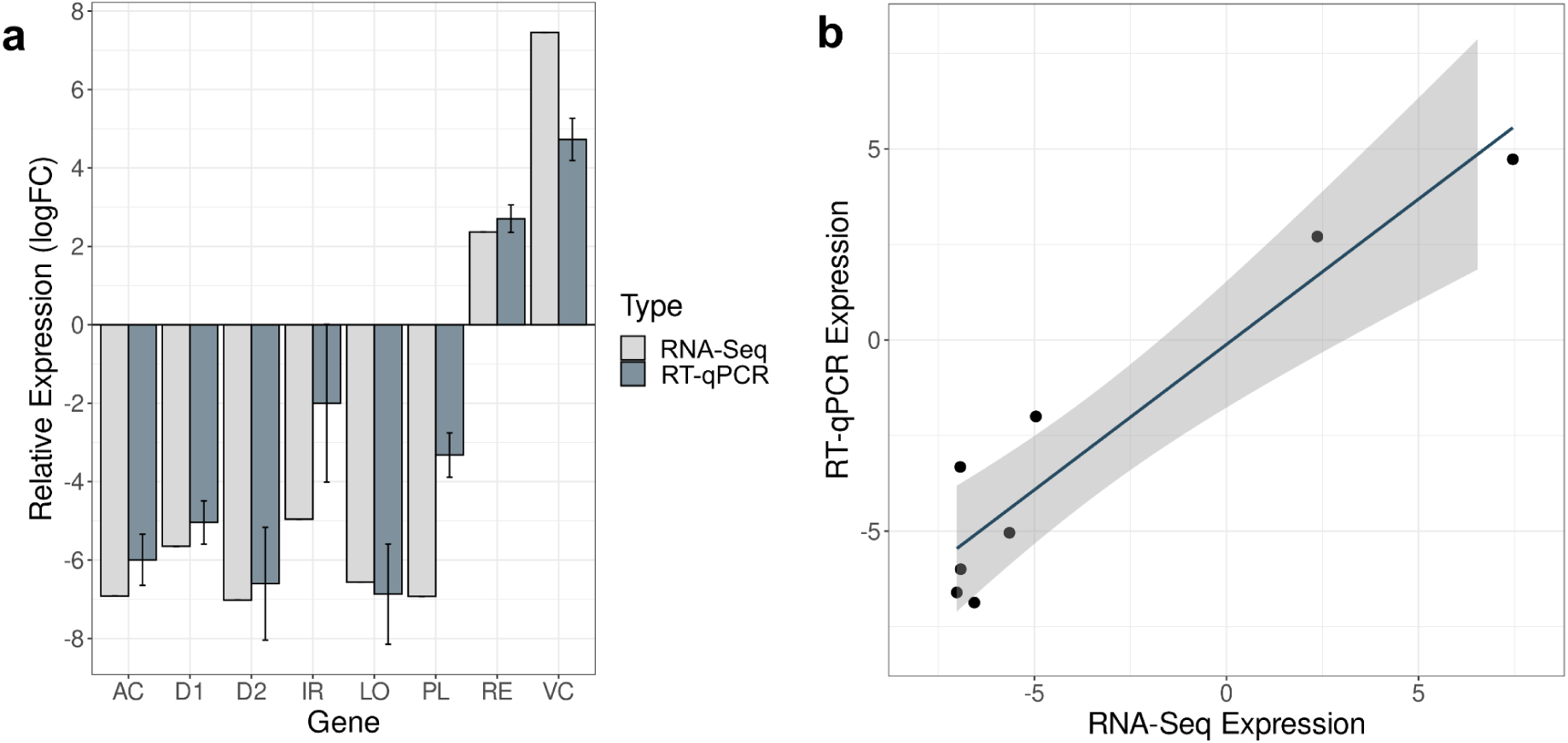
RT‒qPCR transcriptome validation results. (a) Relative expression (log2-fold change) according to RT‒qPCR and transcriptome data are presented as the mean expression value ± SEM (n = 3) relative to that of BGP 344 without infestation (344_C). (b) Correlation between RNA-seq and RT‒qPCR analysis. Abbreviations - AC: acyclic sesquiterpene synthase; D1: unknown transcript 1; D2: unknown transcript 2; IR: protein iron-related transcription; LO: lipoxygenase 2; PL: phospholipase D alpha 2; RE: protein reveille 2; VC: enhancer of mRNA-decapping protein 4.

**Table S2.** Number of total, up- and downregulated DEGs obtained in each treatment comparison.

| DEGs | R0vsR1 | R0vsR2 | R1vsR2 | S0vsS1 | S0vsS2 | S1vsS2 | R0vsS0 | R1vsS1 | R2vsS2 |
| --- | --- | --- | --- | --- | --- | --- | --- | --- | --- |
| Total | 0 | 408 | 30 | 0 | 65 | 41 | 865 | 783 | 1039 |
| Up-regulated | 0 | 1 | 0 | 0 | 14 | 13 | 533 | 469 | 500 |
| Down-regulated | 0 | 407 | 30 | 0 | 51 | 28 | 332 | 314 | 539 |

**Table S3.**
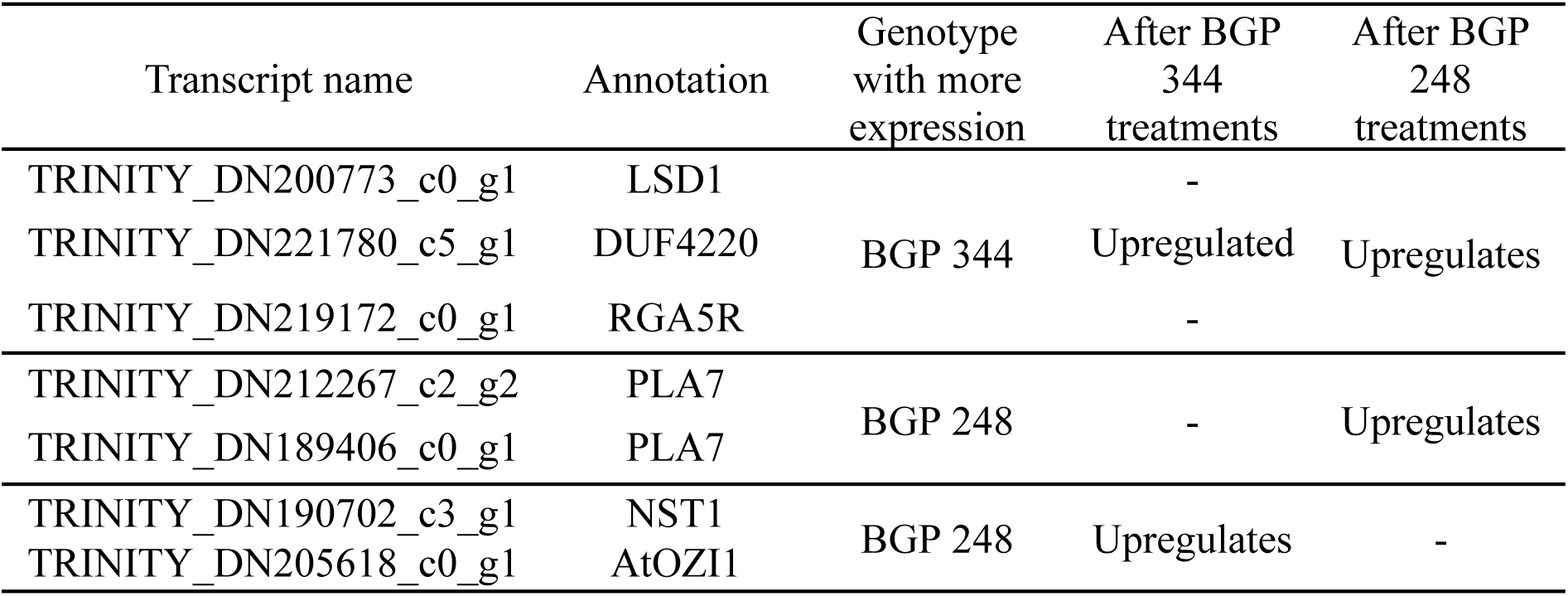
Differentially expressed genes intersecting and their relation to resistance in both genotypes.

**Table S5.** The enriched KEGG pathways are arranged in decreasing *p* value order, and their respective related genotype contrast comparisons are shown.

| KEGG ID | Pathway | Contrast | *p* value |
| --- | --- | --- | --- |
| map00500 | Starch and sucrose metabolism | 344_C vs. 344_2 | 0.044433 |
| map00480 | Glutathione metabolism | 344_1 vs. 248_1 | 0.041352 |
| map00071 | Fatty acid degradation | 248_C vs. 248_2 | 0.038649 |
|  |  | 344_1 vs. 344_2 | 0.025122 |
| map00620 | Pyruvate metabolism | 248_C vs. 248_2 | 0.038649 |
|  |  | 344_1 vs. 344_2 | 0.025122 |
| map04712 | Circadian rhythm - plant | 344_C vs. 248_C | 0.027134 |
| map00592 | alpha-Linolenic acid metabolism | 248_C vs. 248_2 | 0.026678 |
|  |  | 344_1 vs. 344_2 | 0.017209 |
| map00073 | Cutin, suberin and wax biosynthesis | 344_C vs. 248_C | 0.026091 |
| map00010 | Glycolysis/Gluconeogenesis | 248_1 vs. 248_2 | 0.023988 |
|  |  | 248_C vs. 248_2 | 0.018660 |
|  |  | 344_1 vs. 344_2 | 0.009861 |
| map00230 | Purine metabolism | 344_1 vs. 344_2 | 0.017209 |
| map00350 | Tyrosine metabolism | 248_C vs. 248_2 | 0.016575 |
|  |  | 344_1 vs. 344_2 | 0.010610 |
| map00232 | Caffeine metabolism | 344_1 vs. 344_2 | 0.010610 |
| map03030 | DNA replication | 344_C vs. 248_C | 0.010392 |
| map00052 | Galactose metabolism | 344_C vs. 344_2 | 0.000021 |

